# Homeostasis in the Central Dogma of Molecular Biology: the importance of mRNA instability

**DOI:** 10.1101/599050

**Authors:** José E. Pérez-Ortín, Vicente Tordera, Sebastián Chávez

## Abstract

Cell survival requires the control of biomolecule concentration, i.e. biomolecules should approach homeostasis. With information-carrying macromolecules, the particular concentration variation ranges depend on each type: DNA is not buffered, but mRNA and protein concentrations are homeostatically controlled, which leads to the ribostasis and proteostasis concepts. In recent years, we have studied the particular features of mRNA ribostasis and proteostasis in the model organism *S. cerevisiae*. Here we extend this study by comparing published data from three other model organisms: *E. coli, S. pombe* and cultured human cells. We describe how mRNA ribostasis is less strict than proteostasis. A constant ratio appears between the average decay and dilution rates during cell growth for mRNA, but not for proteins. We postulate that this is due to a trade-off between the cost of synthesis and the response capacity. This compromise takes place at the transcription level, but is not possible at the translation level as the high stability of proteins, *versus* that of mRNAs, precludes it. We hypothesize that the middle-place role of mRNA in the *Central Dogma* of Molecular Biology and its chemical instability make it more suitable than proteins for the fast changes needed for gene regulation.

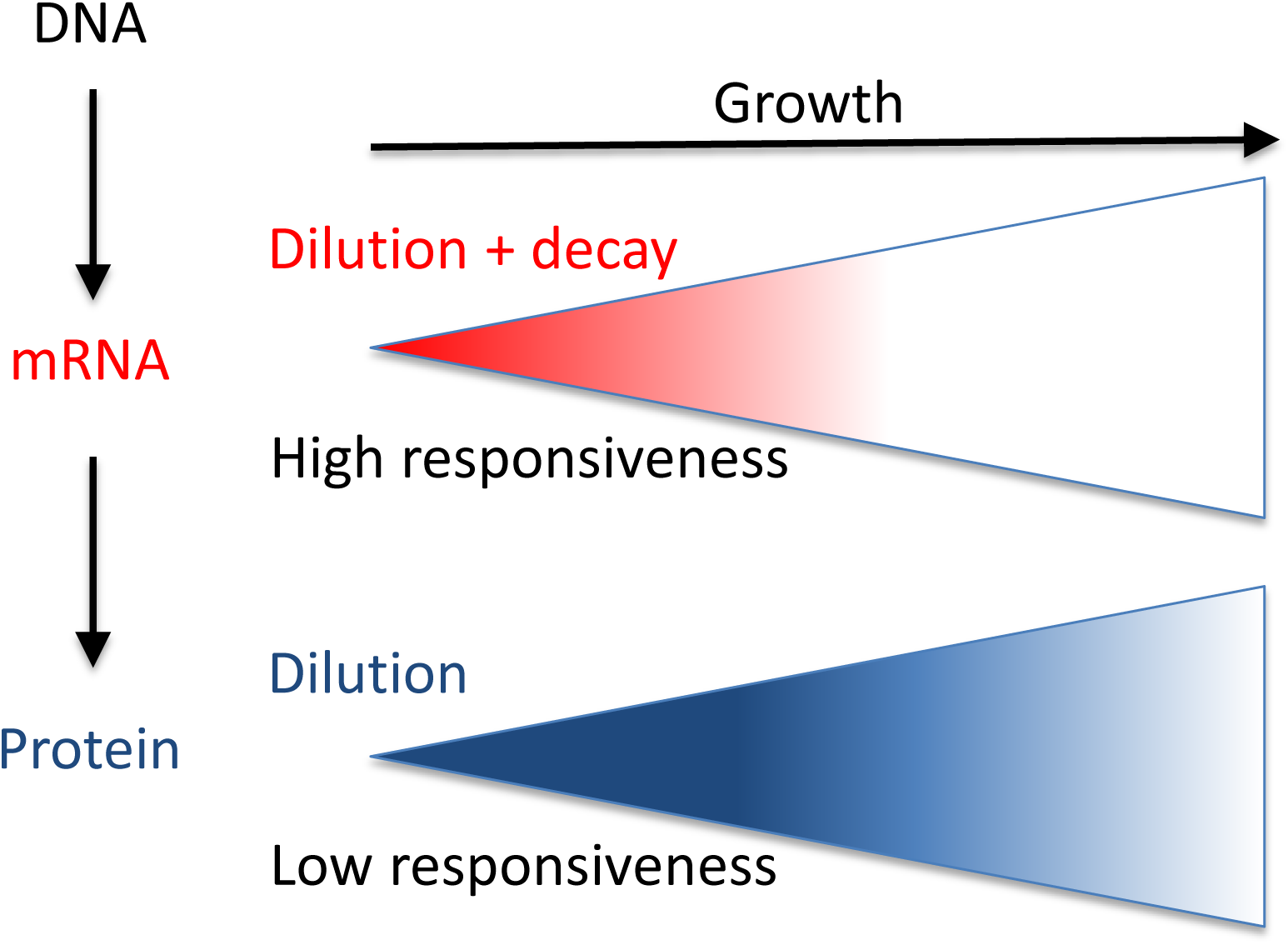
Graphical Abstract.

## Introduction

Homeostasis is the state of steady internal conditions maintained by living things. This concept includes controlling concentrations of cellular molecules and macromolecules. With information-carrying macromolecules, the particular concentration variation ranges depend on molecules. The amount of DNA (genes) cannot be changed along the cell cycle, but in the replication phase. Therefore, it is not quantitatively buffered, but its activity is highly regulated, and its amount determines both nuclei and cell sizes and the transcription process itself (see ref. 29). The importance of the DNA:cell volume ratio has been recently addressed by A. Amon’s group. These authors showed that optimal cell function requires the maintenance of a narrow range of DNA:cytoplasm ratios, and when cell size exceeds the optimal ratio, the cytoplasmic dilution provokes defects in nucleic acids and protein biosynthesis that cause senescence in both yeasts and human cells (37).

On the contrary, mRNAs and proteins have no fixed amount per cell. They are multiple independent molecules that vary upon demand. A fundamental idea lies behind gene regulation: genes numbers are fixed, but their expression can change. Gene expression follows an obligatory flux called the *Central Dogma* (12). As the concentrations of both individual species vary, it is possible to think that homeostatic control for mRNAs and proteins is not necessary. However, the total protein concentration in all cells is known to remain quite constant (24,35). The total mRNA concentration has not been very well studied, and is generally assumed to vary within a certain range; i.e. to be homeostatically controlled (5,42). The terms ribostasis and proteostasis are commonly used to respectively refer to RNA (normally meaning mRNA) and protein homeostasis.

In order to understand the particular features of mRNA ribostasis and proteostasis, we have studied variations in the total mRNA and protein concentrations ([mRNA], [protein]) between different growth conditions in the yeast S*accharomyces cerevisiae* over the last decade. We have also studied the mechanisms to keep homeostasis by means of transcription and degradation rates, and their cross-talk mechanisms (5, 15, 17, 30). Another approach to understand the different roles of mRNAs and proteins, and their various homeostasis features, is to compare total [mRNA] and [protein], and those of their synthesis machineries, among the model organisms ranging from bacteria to human cells. Here we review published data from our own group and other groups on four model organisms for which enough information is available: *Escherichia coli, S. cerevisiae, Schizo*s*accharomyces pombe* and human cells in culture related to [mRNA] and [protein], synthesis and stabilities (half-lives), and also for their synthesis machineries: RNA polymerases (RNA pol) and ribosomes. With these data, we have compared organisms, and also all these parameters, to extract the general rules for the mRNA and protein homeostasis in them. A recent paper by Hauser et al. (18) followed a similar strategy by comparing omics data from four model organisms: *E. coli, S. cerevisiae*, cultured mouse and human cells. That study, however, focused only on the synthesis rates of mRNAs and proteins, and found biological noise (46) to be a key element in the selection of regulatory expression strategies for each protein-coding gene. In this study, however, we focus on how global mRNA and protein concentrations are maintained throughout evolution by considering both synthesis and decay rates.

Using all these quantitative data, we conclude that the evolution of the gene expression flux (Central Dogma) and the chemical properties of RNAs and proteins can explain how mRNA became the main point of gene regulation because of its instability and the central position of the flux. This regulatory role of mRNA made transcription the main point for biological noise. mRNA ribostasis is, therefore, not as strict as proteostasis because of the need for mRNA changes and the comparatively low cost of mRNA transcription.

## Materials and Methods

### Calculating transcription rates for eukaryotic RNA polymerases

Transcription rates (TR) are usually defined based on rNTP consumption. In *S. cerevisiae*, they were calculated as 60%, 25% & 15% of the total consumption for RNA pol I, II & III, respectively (59). Another way to define the relative importance of each RNA pol is regarding the production rate of transcripts that each one synthetizes. In the same yeast, the number of produced rRNAs (ribosomes) has been estimated at 300000 per cell cycle of 90 min (58) or 2000/min (59), e.g. RNA pol I transcribes between 33-55 35S rRNA transcripts/s. As 5S rRNA should be coordinated with 35S rRNA, we assume that its synthesis rate should be the equivalent (≈45 transcripts/s). tRNAs have been estimated to be produced at 3000000 per cell cycle (58) or about 555 transcripts/s. Thus RNA pol III transcribes about 600 molecules/s (Figure 1). Synthesis of mRNAs by RNA pol II can be calculated from our previous study (41), but after correcting for the newest data of mRNAs abundances and stabilities (Tables I & II) at about 60 mRNAs/s. Therefore regarding the transcript number production, the respective contributions of RNA pol I, II and III are 6.5%, 8.5% and 85%, respectively. These percentages can be recalculated using rNTP consumption data (see above) and taking into account the average sizes of their transcripts. RNA pol III transcribes much shorter molecules (0.13 kb on average for 5S and tRNAs primary transcripts) than RNA pol II (1.5 kb mRNA on average, ref. 13) and RNA pol I (6.9 kb 35S transcript). The results are offered in the following percentages about transcribed molecules: 6%, 12% and 82% respectively for RNA pol I, II and III (Figure 1). These figures well fit the previous calculations for the transcribed copy numbers of the three RNA pols. Finally as for the number of different transcribed genes, budding yeast RNA pol II transcribes most genes, i.e. >90%, because there are about 6000 protein-encoding genes *vs*. ≈300 RNA pol III genes, and only one RNA pol I gene (with 100-200 copies).

**Table I:** Abundances of molecules and synthesis machineries. Data refer to the cells growing at their highest growth rate. They have been obtained from references 2,4,7,16 and 33-35 for *E. coli*; 19,35,41,56,58 and 62 for *S. cerevisiae*; 28,29 and 51 for *S. pombe* and 8,35,48 50, 55 for HeLa and other mammalian cells. The data for RNA polymerases I, II & III were calculated from the proteomic datasets of *S. cerevisiae, S. pombe* and HeLa (19,21,40) using an average of the three largest specific subunits of each. *The data for *S. pombe* and HeLa were corrected as described by Milo (33). No data for RNA pol III in *S. pombe* and RNA pol I & III in HeLa have been found in the datasets. Cell sizes correspond to whole cell volume and were taken from references 16, 33, 35 & 51.

|  | *S. cerevisiae* | *S. pombe* | HeLa | *E. coli* |
| --- | --- | --- | --- | --- |
| mRNAs molecules /cell | $3-6 \cdot 10^4$ | $1-2 \cdot 10^5$ | $3-4 \cdot 10^5$ | $8 \cdot 10^3$ |
| Proteins: molecules /cell | $8 \cdot 10^7$ | $3 \cdot 10^8$ | $5 \cdot 10^9$ | $4-8 \cdot 10^6$ |
| rRNA (ribosomes): molecules /cell | $2-3 \cdot 10^5$ | $5 \cdot 10^5$ | $4-10 \cdot 10^6$ | $5-7 \cdot 10^4$ |
| tRNAs: molecules /cell | $3 \cdot 10^6$ | ? | $3 \cdot 10^7$ | $4-7 \cdot 10^5$ |
| Cell size (fL) | 50 | 150 | $3 \cdot 10^3$ | 1-3 |
| [mRNA: molecules/fL] | $1 \cdot 10^3$ | $1 \cdot 10^3$ | $1 \cdot 10^2$ | $4 \cdot 10^3$ |
| [protein]: molecules/fL | $2 \cdot 10^6$ | $1-2 \cdot 10^6$ * | $1-2 \cdot 10^6$ * | $3 \cdot 10^6$ |
| [ribosomes]: molecules/fL | $4-7 \cdot 10^3$ | $4-5 \cdot 10^3$ | $1-4 \cdot 10^3$ | $3 \cdot 10^4$ |
| RNA pol II : molecules /cell | $2-3 \cdot 10^4$ | $2 \cdot 10^4$ * | $2-4 \cdot 10^5$ | $11 \cdot 10^3$ |
| RNA pol I: molecules /cell | $8-15 \cdot 10^3$ | $2 \cdot 10^4$ * | ? | |
| RNA pol III | $5 \cdot 10^3$ | ? | ? | |
| RNA pol II /ribosomes | 0.1-0.2 | 0.04 | 0.03-0.05 | 0.1 |
| RNA pol I /ribosomes | 0.05-0.2 | 0.04 | ? |  |
| Ribosomes/mRNA | 2-11 | 3-4 | 9-30 | 15-30 |

**Table II:** Generation time (GT) correlations with ribosome and mRNA concentration and mRNA and protein stabilities. Data refer to the cells growing at their highest growth rate (minimal generation time, GT) and were obtained from references 27,36 for *E. coli*; 39,44,51 for *S. cerevisiae*; 51 for *S. pombe* and 48, 53 for HeLa and other cultured mammalian cells. Note that half-lives (HL) are in minutes for mRNAs and in hours for proteins.

|  | GT (min) | mRNA<br>HL<br>(min) | mRNA<br>HL/GT | protein<br>HL (hr) | Protein<br>HL/GT | [Ribosome]<br>x GT | [mRNA]<br>relative | [mRNA]<br>x GT |
| --- | --- | --- | --- | --- | --- | --- | --- | --- |
| *E. coli* | 20 | 1-5 | 0.13 | 5-20 | >15 | $4 \cdot 10^5$ | 1 | 20 |
| *S. cerevisiae* | 90 | 8-15 | 0.15 | 8.8 | 6 | $4\text{-}6 \cdot 10^5$ | 0.35 | 30 |
| *S. pombe* | 100 | 15-40 | 0.28 | 12 | 7 | $4 \cdot 10^5$ | 0.35 | 30 |
| HeLa | $1.5\text{-}3 \cdot 10^3$ | 290-600 | 0.1-0.2 | 36 | 0.7-1.5 | $2\text{-}8 \cdot 10^6$ | 0.035 | 50 |

**Figure 1.**
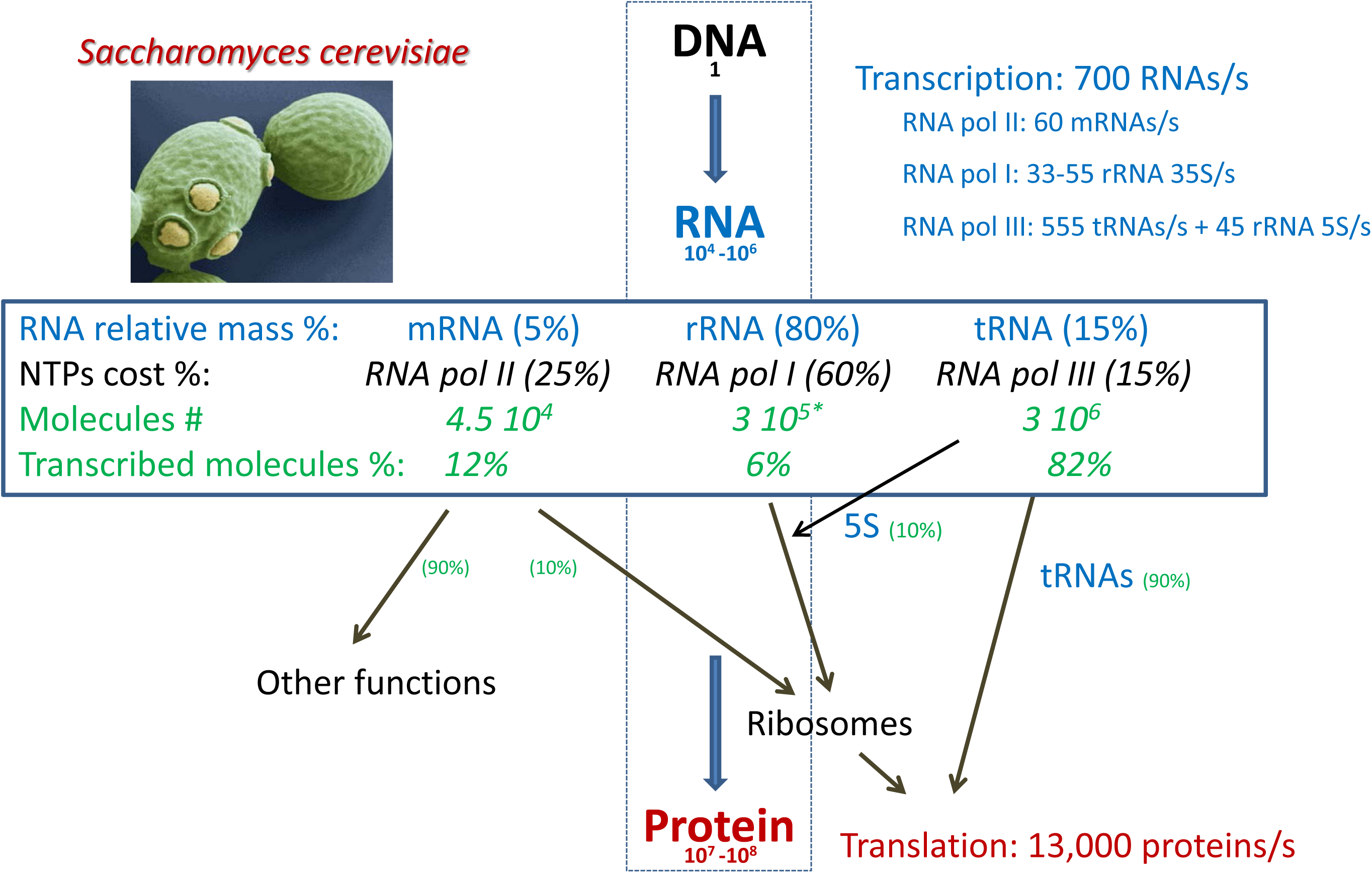
Quantitation of the Central Dogma in eukaryotes. Using data from published references (see M&M for a detailed description), we herein represent the amounts of the molecules involved in the flux of information from DNA (genome) to protein in *S. cerevisiae* as an example of eukaryotic cells. It should be similar, in relative proportions, for other eukaryotes. For a haploid budding yeast, the number of RNAs falls within the range of 10^−4^−10^−6^, including mRNAs, tRNAs and rRNAs. Other RNAs come in much smaller amounts and are not considered herein. The relative proportions of RNA types and transcription costs are shown (see M&M for details). The percentage of RNA pol II transcription devoted to ribosome-related proteins (10%) and the other proteins (from ref. 41) and that of RNA pol III devoted to tRNAs and 5S rRNA (see M&M) are shown. Note that as the total number of protein molecules and their total translation rates are higher than their mRNA counterparts, the translation cost is much higher. The * indicates that we here take the number of ribosomes as number or molecules of rRNA, but it should be considered that each ribosome has 4 independent rRNA molecules, three transcribed by RNA pol I as a single nascent transcript: 5.8S, 18S and 25S; and another transcribed by RNA pol III: 5 S.

## Results

This paper reviews published quantitative data from RNAs and proteins in four model organisms to conclude that there are universal rules for the proportions of both molecules and their synthesis machineries, except for the very variable total [mRNA]. Thus we postulate that the poor stability of mRNAs and the relatively good stability of proteins influence their cellular functions, and explain why most gene regulation occurs at the transcriptional level. Finally, we discuss these differences between mRNAs and proteins from an evolutionary point of view.

### Why proteostasis is much stricter than mRNA ribostasis

We collected published data about the abundances of proteins and mRNAs molecules in three free-living microorganisms: the eubacteria *E. coli*, and two distantly related yeasts *S. cerevisiae* and *S. pombe*, and from human cells in culture (mainly HeLa cells for most data). Table I shows that mRNAs and proteins have very different average numbers in all organisms. It should be noted that the data obtained from different laboratories vary by one order of magnitude. For this reason, we herein used data that are supported by more than one study or, alternatively, we show the range of published values. It is also important to consider that we are talking about total populations of molecules of many different types with considerable variations between them, and that do not follow normal distributions. In any case, the differences between proteins and mRNAs are robust enough to allow differential conclusions to be made. Proteins are thousands of times more abundant, but have variable factors depending on the organism: about a 10^3^ factor in microorganisms and one of 10^4^ in human cells. The number of each class of molecules scales with cell size. The scale is quite constant for proteins, which remain [protein] uniform (1-3 million molecules/fL), as formerly stated by Milo et al (33–34). However, the [mRNA] is more variable, about 4-fold higher in *E. coli* than in yeasts and 10-fold lower in human cells. Whether this major change reflects a functional property (see below) remains an open question. The large number of protein molecules is probably the reason for the conservation of the total [protein] between different living beings, and also between distinct physiological situations (strict proteostasis). This is because cell mass is composed a large proportion of protein (about 50% of dry weight in most organisms: 22,34). In contrast, total RNA is much less abundant: 4-10% in the dry weight mass in budding yeast (22), and 15-20% in *E. coli* (47,52). Moreover, most RNA is not mRNA, which constitutes only a small fraction (5-10% of the total RNA in all organisms: 26,39,52,59). So the fraction of cell mass that is mRNA is so tiny (1% at the most) that mRNA ribostasis cannot impact the structural features of cells, unlike proteostasis.

The abundance of proteins and mRNAs should, on the other hand, require a corresponding abundance of their synthetic machineries (RNA polymerases and ribosomes). Ribosomes are much more abundant than RNA pol (5-33x, Table I), but this ratio is much lower than the relative amounts of proteins and mRNAs (>1000x). This difference indicates that synthesis rates are not the only factor to explain the actual difference in quantities: the differential stabilities of mRNAs and proteins also matter (see below). On the other hand, ribosomes have similar concentration in eukaryotes (about 4000/fL), but are 5-fold more concentrated in *E. coli*. This can be caused by the higher concentration of protein molecules, by the shorter generation time of this bacterium and/or by the characteristic coupling between transcription and translation in prokaryotes.

When putting together the quantitative data for molecules and their synthesis rates in the Central Dogma, several interesting features appear. Figure 1 summarizes the data for budding yeast. The data known for other eukaryotes are quite similar (in relative terms), so we consider that this figure can act as a general profile for a eukaryotic cell. A single DNA molecule (haploid genome) produces RNAs that are in the steady state within the range from 10^4^ to 10^6^ total molecules per cell (less than 10^5^ for mRNAs; see Table I). Proteins are much bigger in number (10^7^ to 10^8^), which affects the cost of translation. By using published data, we calculate that the total TR is about 700 RNAs/second in *S. cerevisiae*. This synthesis rate is split into three RNA polymerases (see M&M) because transcription is not only devoted to make mRNAs. Other ncRNAs are also transcribed and perform important functions. If we set a limit to the most abundant ncRNAs, those that participate in the translation process, e.g. tRNAs and rRNAs, then their transcription in eukaryotes is done by two specialized RNA pol: I and III. They are only slightly less abundant than RNA pol II (Table I) in spite of transcribing much fewer genes because they have to produce a 10-fold larger number of transcripts (Fig. 1). In fact each RNA pol type plays a maximal quantitative role depending on the analyzed parameter. RNA pol I consumes the most rNTPs, RNA pol III is the biggest producer of transcripts and RNA pol II has the most target genes (see Figure 1 and M&M).

In quantitative terms, protein synthesis is, however, much higher than mRNA synthesis: 13000 (56) *vs*. 700 molecules/s. The obvious consequence is that translation is much more costly than total transcription, and even more so if only mRNA transcription by RNA pol II is considered. The energy cost of total translation in *S. cerevisiae* has been calculated to be about 10-20x higher than mRNA transcription (57). This author used old median mRNA stability (23 min) and protein stability (30 h) values. If the most recent and confident data were used (see Table II), mRNA translation into proteins should be even more costly (30x or more) than gene transcription. Thus, energy cost is a key parameter that should determine the strategies of cells for [mRNA] and [protein] homeostasis.

### What the numbers of molecules and their turnover rates can tell us about gene expression

mRNAs and proteins also have very different stabilities in all living cells (Table II). mRNA half-lives scale with the generation time (GT) between different cells, and also between the GTs of a single species. The mRNAs median half-life is usually about 15-20% of the cell’s GT (Table II). This has already been pointed by other authors (35,53,61). Given the much higher GT than the median mRNA half-life, transcription by RNA pol II is devoted mainly to compensate mRNA degradation (41). We previously proposed that the approximately constant mRNA half-life /GT ratio could be due to the need for an optimal balance to be maintained between the synthesis cost and the response capacity (9). Here we propose that this might be a general rule for all cells growing at their highest growth rate (Figure 2).

**Figure 2.**
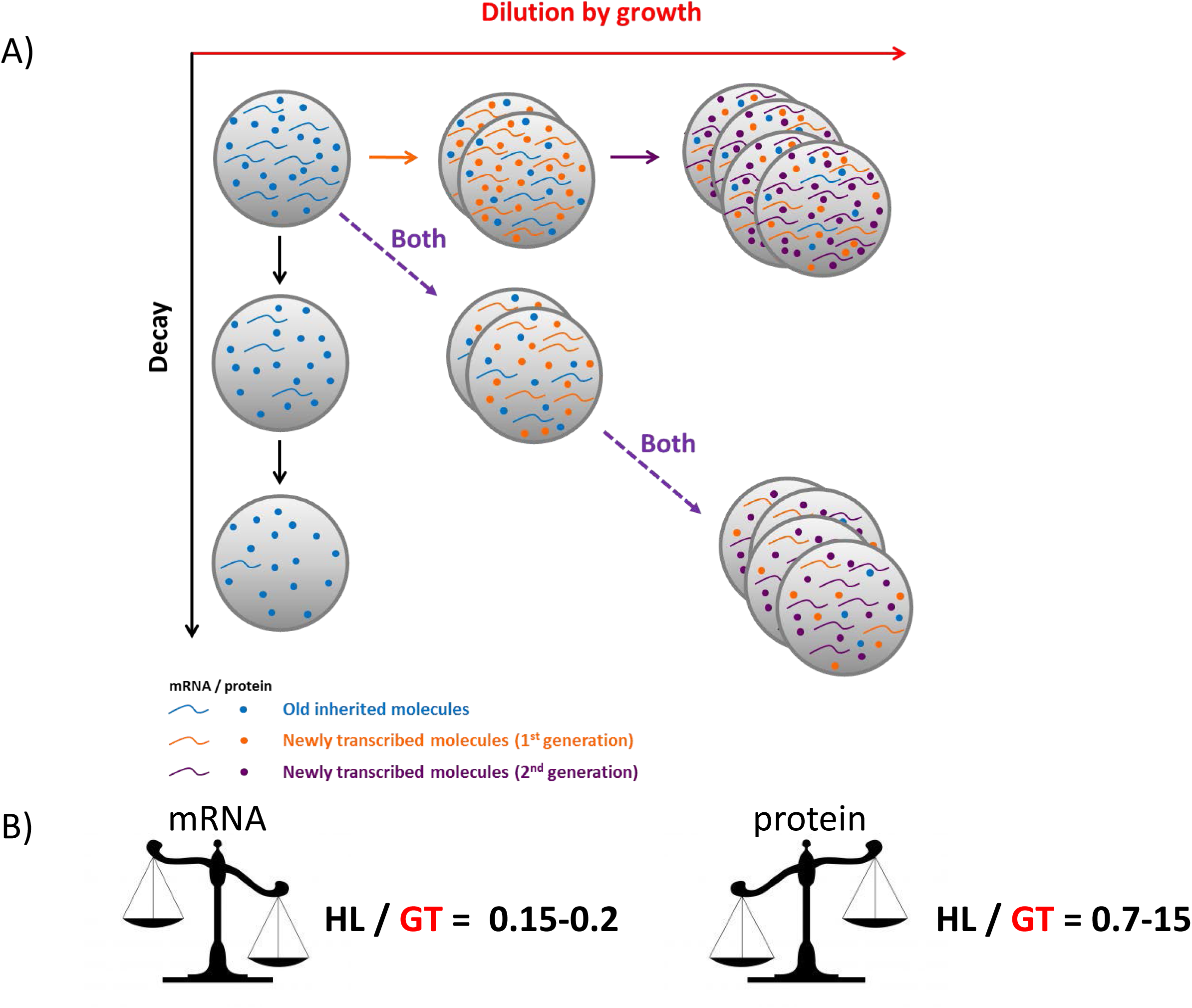
A comparison of the relative contributions of the decay and dilution rates for mRNAs and proteins. A) In actively dividing cells, the synthesis rates for any molecule should compensate both the degradation (decay, in y axes) and dilution caused by an increase in cell mass/volume (growth, x axes) if a homeostatic concentration is to be maintained (both processes, diagonal). Note that horizontal and vertical represent the hypothetical situations with only one process occurring. B) For mRNAs, which are relatively short-lived molecules, the proportion of transcription devoted to compensate the degraded molecules and dilution, expressed herein as the ratio between half-lives (HLs) and GTs, is approximately constant for all the studied organisms: about 15-20%. We interpret this as a trade-off between response capacity and synthesis costs. On the contrary, for those proteins that are very stable and more abundant than mRNAs, synthesis is devoted mostly to compensate dilution with a HL/GT ratio that is very high and variable. Thus for proteins, a trade-off between response capacity and synthesis costs does not seem to be an evolutionary constraint (see the main text for details).

Protein half-lives do not, however, scale with the GT. The protein median half-life/GT ratio varies among organisms (Table II). Protein half-lives in microorganisms are much longer than GTs, and take a similar value in human cells in culture. Thus most translation seems devoted to compensate the dilution caused more by growth rather than by degradation (10,60), except in cultured human cells in which the contributions of both are similar (Table II). It should be noted that in nondividing cells, translation is quantitatively much less important because only protein degradation should be compensated (63). Hence in non-dividing cells, the global transcription/translation cost balance differs considerably from actively growing cells.

So although the response capacity of cells to environmental changes is related to proteins because they are the final goal of gene expression, and even when considering that a cell’s translation cost is much higher than the mRNA transcription cost (Figure 1), there is no single optimal balance between synthesis cost and response capacity in proteins for all organisms. This balance depends on the particular type of organism, bacteria, yeasts or mammalian cells, and on the particular GT. The cost of making proteins is high, but is not affected much by their half-lives because degradation is only a small part of protein disappearance (especially in free-living microorganisms), unlike mRNA (Figure 2). This can be due ultimately to the intrinsically higher chemical stability of proteins and should have been a key factor during primitive life evolution, which selected protein-based cells *vs*. RNA-based ones. Therefore, the intrinsic (and very convenient for living cells) high stability of proteins has precluded an evolutionary search for an optimal balance between synthesis cost and response capacity. Response capacity in proteins is, therefore, based mostly on post-translational modifications not on translational regulation (23)

We conclude that the marked strictness of global proteostasis and the high stability of most proteins, compared to mRNAs, explain why evolution has selected mainly regulatory mechanisms at the mRNA level for most genes.

### Variation in mRNA concentration is related to translation rate

Finally, another question that arises from the quantitative data in Table I is the relative numbers of the three components in the translation process. The ribosomes/mRNAs ratio tends to be similar in microorganisms, some 4-10 ribosomes/mRNA, which is not far from the *in vivo* experimental ribosome densities measured in *S. cerevisiae* and *E. coli* (6-7 ribosomes/mRNA, ref. 1, 6). This is true even when the [ribosome] decreases because of the different growth temperature (5), which suggests that it is mechanistically constrained. In fact, it has been shown in both *E. coli* and *S. cerevisiae* that, when grown at their fastest growth rate (i.e. the shortest GT), ribosomes are saturated with mRNAs (11). It is likely that ribosomes are almost saturated with mRNAs in all microorganisms when grown at their fastest rate. Thus if we multiply [ribosome] x GT, an almost constant value is obtained (Table II). This is explained by protein production being limited by the availability of free ribosomes (49), and also explains why the GT is limited by protein production as proteins represent a high percentage of cell mass (see above). It has been shown in both *E. coli* and *S. cerevisiae* that [ribosomes] depend on the GT within a range of fast growth conditions. In *E. coli*, the percentage of protein devoted to ribosomes (ribosome/dry mass) grows at a growth rate that is almost proportionally at a GT between 60 and 24 min (35). In the *S. cerevisiae* grown in different media (57), ribosome x GT is also constant for a wide range of GTs (from 240 to 90 min). In both cases this constant parameter is not maintained at a longer GT, where excess ribosomes per mRNA seem to occur, similarly to what happens in human cells (54 & Table 1). In cultured human cells, the [ribosome] x GT is much higher, which suggests different constraints for translation in multicellular organisms. On the other hand, in *E. coli* and *S. cerevisiae*, it has been experimentally demonstrated that certain excess ribosomes appear even at the fastest growth rate, and this fraction increases as the growth rate decreases. It has been suggested that these excess ribosomes are employed when translation unexpectedly demands an increase (20,31).

Interestingly as mentioned before, [mRNA] varies more widely between organisms than [protein] (Table I). However, and strikingly, [mRNA] approximately correlates inversely to the GT (the product of both is almost constant) when comparing the four organisms (Table II). This is also seen when comparing a single organism at very different GTs. For instance, it has been shown that [mRNA] decreases after a diauxic shift when the GT increases (45) and translation decreases. Thus we propose that, at least for microorganisms, wide variations in [mRNA] are related to translation rate control as the ribosomes/mRNA ratio is constant and the [ribosome] determines the maximum translation rate capacity. To support this hypothesis, it has been recently suggested that the total [mRNA] in *S. cerevisiae* influences the GT because it defines the total number of ribosomes engaged in translation (32). This would explain that both [ribosome] and [mRNA] change in parallel when growing budding yeasts at different growth temperatures (5).

### The role of biological noise

As pointed out in the Introduction, Hausser et al (18) compared the abundances and synthesis rates for individual genes in four model organisms. [mRNA] steady-state levels are the result of mRNA synthesis and degradation rates. Although these authors did not analyze the latter in detail, they concluded that biological noise is a key element in the selection of regulatory expression strategies for protein-coding genes. It is known that intrinsic noise is due mainly to transcription, and not to translation (38,46), which is owing chiefly to fluctuations in the mRNA levels that arise from the stochastic activation of gene promoters (3). They argue that as the intrinsic noise for two proteins with identical abundance is higher for those with lower [mRNA] (e.g. less TR because mRNA stability does not strongly influence [mRNA]), cells can choose an expression strategy with high noise (low TR) or low noise (high TR) levels. This is not strictly true because the genes with the same [mRNA] can have different TR due to their varying stabilities, and this is an important expression strategy for a gene (43). In fact it has been suggested that both the birth and death of mRNAs can affect noise (3). In any case, a higher TR means a higher cost. Although the transcription cost is much lower than the translation cost (Fig. 1), it is important in evolutionary terms because the translation cost is fixed for a given protein with a given abundance to play its biological role. The transcription cost is, however, elective for every gene by it striking a balance between precision and economy depending on TR.

The history of RNA, its place in the middle of the expression flux and its chemical properties should have been the origins for selecting transcriptional regulation, including epigenetic mechanisms, instead of translational regulation. Therefore, the equilibrium between the noise and cost caused by the election of a given TR should have been a crucial factor to select an expression strategy for each gene.

### Conclusion: a historical and functional perspective of the Central Dogma

In the RNA world, early life used RNA for both genetic information and catalytic ability (14). RNA is more unstable and mutable than DNA and proteins (25). Therefore, RNA high turnover should always have been a key factor of its function. When proteins appeared in primitive cells, they substituted RNAs for most structural and catalytic functions because of their versatility and stability. After DNA appeared later (14), RNA was positioned as an obligate intermediate in the gene expression flux (Central Dogma) and became the main point of gene regulation given its instability and central position (mRNA) in the flux. A theoretical question is why mRNA remains in the middle and was not replaced completely with DNA for information storage, and with protein for catalytic functions, to make life with a simpler Central Dogma (DNA→protein). One possible answer to this question is that cells have taken advantage of the natural instability of mRNA, instead of protecting it as they did for other stable RNAs (rRNA, tRNA, etc.), because it allows better and more flexible regulatory mechanisms. Although the mRNA amount can be regulated at both synthesis (TR) and degradation rates (half-life) (43), there is a strong preference for regulation at the synthesis level (23). This also made the TR the main point for biological noise.

In conclusion, mRNA ribostasis is less strict than proteostasis because of the regulatory need for mRNA changes and the comparatively lower cost of mRNA synthesis. This causes a trade-off between cost and response capacity at the transcription level. The high protein stability, and also the higher protein synthesis cost being much higher than the gene transcription cost, explain why protein turnover has not been evolutionary conformed as a general way to regulate gene expression.

## Acknowledgements

We thank Drs. José García-Martínez, A. Jordán-Pla and Vicent Pelechano for their helpful discussions. This work has been supported by grants from the Spanish Ministry of Economy and Competitiveness, and European Union funds (FEDER) [BFU2016-77728-C3-1-P to S. C.], [BFU2016-77728-C3-3-P and BFU2015-71978-REDT to J.E.P-O], from the Regional Valencian Government [PROMETEO II 2015/006 to J.E.P-O].

## References

1. Arava Y, Wang Y, Storey JD, Liu CL, Brown PO, Herschlag D. 2003. Genome-wide analysis of mRNA translation profiles in Saccharomyces cerevisiae. Proc Natl Acad Sci U S A. 100(7):3889–3894.

2. Bakshi S, Siryaporn A, Goulian M, Weisshaar JC. 2012. Superresolution imaging of ribosomes and RNA polymerase in live Escherichia coli cells. Mol Microbiol. 85(1):21–38. doi: 10.1111/j.1365-2958.2012.08081.

3. Bar-Even A, Paulsson J, Maheshri N, Carmi M, O’Shea E, Pilpel Y, Barkai N. 2006. Noise in protein expression scales with natural protein abundance. Nat Genet. 38(6):636–43.

4. Bartholomäus A et al., 2016. Bacteria differently regulate mRNA abundance to specifically respond to various stresses. Philos Trans A Math Phys Eng Sci. 374(2063). pii: 20150069. doi: 10.1098/rsta.2015.0069

5. Benet M, Miguel A, Carrasco F, Li T, Planells J, Alepuz P, Tordera V, Pérez-Ortín JE. 2017. Modulation of protein synthesis and degradation maintains proteostasis during yeast growth at different temperatures. Biochim Biophys Acta Gene Regul Mech. 1860(7):794–802. doi:10.1016/j.bbagrm.2017.04.003.

6. Brandt F, Etchells SA, Ortiz JO, Elcock AH, Hartl FU, Baumeister W. 2009. The native 3D organization of bacterial polysomes. Cell 136(2):261–271.

7. Bremer, H., Dennis, P. P. 1996. Modulation of chemical composition and other parameters of the cell by growth rate. Neidhardt, et al. eds. Escherichia coli and Salmonella typhimurium: Cellular and Molecular Biology, 2nd ed. chapter 97, pp. 1559.

8. Carter MG, Sharov AA, VanBuren V, Dudekula DB, Carmack CE, et al. 2005. Transcript copy number estimation using a mouse whole-genome oligonucleotide microarray. Genome Biol 6: R61

9. Chávez S, García-Martínez J, Delgado-Ramos L, Pérez-Ortín JE. 2016. The importance of controlling mRNA turnover during cell proliferation. Curr Genet. 62(4):701–710.

10. Christiano R, Nagaraj N, Fröhlich F, Walther TC. 2014. Global proteome turnover analyses of the Yeasts S. cerevisiae and S. pombe. Cell Rep. 9(5):1959–1965. doi: 10.1016/j.celrep.2014.10.065.

11. Chu D, von der Haar T. 2012. The architecture of eukaryotic translation. Nucleic Acids Res. 40(20):10098–106. doi: 10.1093/nar/gks825.

12. Crick FH. 1958. On protein synthesis. Symp Soc Exp Biol. 12:138–163.

13. Dujon B. 1996. The yeast genome project: what did we learn? Trends Genet. 12(7):263–270.

14. Dworkin JP, Lazcano A, Miller SL. 2003. The roads to and from the RNA world. J Theor Biol. 222(1):127–134.

15. García-Martínez J, Delgado-Ramos L, Ayala G, Pelechano V, Medina DA, Carrasco F, González R, Andrés-León E, Steinmetz L, Warringer J, Chávez S, Pérez-Ortín JE. 2016. The cellular growth rate controls overall mRNA turnover, and modulates either transcription or degradation rates of particular gene regulons. Nucleic Acids Res. 44(8):3643–3658. doi: 10.1093/nar/gkv1512.

16. Grossman N, Ron EZ, Woldringh CL. 1982. Changes in cell dimensions during amino acid starvation of Escherichia coli. J Bacteriol. 152(1):35–41

17. Haimovich G, Medina DA, Causse SZ, Garber M, Millán-Zambrano G, Barkai O, Chávez S, Pérez-Ortín JE, Darzacq X, Choder M. 2013. Gene expression is circular: factors for mRNA degradation also foster mRNA synthesis. Cell 153(5):1000–1111. doi: 10.1016/j.cell.2013.05.012.

18. Hausser J, Mayo A, Keren L, Alon U. 2019. Central dogma rates and the trade-off between precision and economy in gene expression. Nat Commun. 10(1):68. doi: 10.1038/s41467-018-07391-8.

19. Ho B, Baryshnikova A, Brown GW. 2018. Unification of Protein Abundance Datasets Yields a Quantitative Saccharomyces cerevisiae Proteome. Cell Syst. 6(2):192–205.e3. doi: 10.1016/j.cels.2017.12.004

20. Kafri M, Metzl-Raz E, Jona G, Barkai N. 2016. The Cost of Protein Production. Cell Rep. 14(1):22–31. doi: 10.1016/j.celrep.2015.12.015.

21. Kulak NA, Pichler G, Paron I, Nagaraj N, Mann M. 2014. Minimal, encapsulated proteomic-sample processing applied to copy-number estimation in eukaryotic cells. Nat Methods. 11(3):319–24. doi: 10.1038/nmeth.2834.

22. Lange HC, Heijnen JJ. 2001. Statistical reconciliation of the elemental and molecular biomass composition of Saccharomyces cerevisiae. Biotechnol Bioeng. 75(3):334–44.

23. Li JJ, Biggin MD. 2015. Gene expression. Statistics requantitates the central dogma. Science 347(6226):1066–7. doi: 10.1126/science.aaa8332.

24. Liebermeister W, Noor E, Flamholz A, Davidi D, Bernhardt J, Milo R. 2014. Visual account of protein investment in cellular functions. Proc Natl Acad Sci U S A. 111(23):8488–8493. doi:10.1073/pnas.1314810111.

25. Lindahl, T. (1993). Instability and decay of the primary structure of DNA. Nature 362: 709–715.

26. Lodish H, et al. (2003). Molecular Cell Biology. 5th edition W. H. Freeman & Co Ltd. ISBN-10: 0716743663.

27. Mackie GA. 2013. RNase E: at the interface of bacterial RNA processing and decay. Nat Rev Microbiol. 11(1):45–57. doi: 10.1038/nrmicro2930.

28. Maclean N. 1965. Ribosome numbers in a fission yeast. Nature. 207:322–323.

29. Marguerat S, Schmidt A, Codlin S, Chen W, Aebersold R, Bähler J. 2012. Quantitative analysis of fission yeast transcriptomes and proteomes in proliferating and quiescent cells. Cell 151(3):671–83. doi: 10.1016/j.cell.2012.09.019.

30. Mena A, Medina DA, García-Martínez J, Begley V, Singh A, Chávez S, Muñoz-Centeno MC, Pérez-Ortín JE. 2017. Asymmetric cell division requires specific mechanisms for adjusting global transcription. Nucleic Acids Res. 45(21):12401–12412. doi: 10.1093/nar/gkx974.

31. Metzl-Raz E, Kafri M, Yaakov G, Soifer I, Gurvich Y, Barkai N. 2017. Principles of cellular resource allocation revealed by condition-dependent proteome profiling. Elife. 6. pii: e28034. doi: 10.7554/eLife.28034.

32. Metzl-Raz E, Kafri M, Yaakov G, Barkai N. (2019). Gene transcription as a limiting factor in protein production and cell growth. BioRxiv doi: http://dx.doi.org/10.1101/626242.

33. Milo R. 2013. What is the total number of protein molecules per cell volume? A call to rethink some published values. Bioessays 35(12):1050–1055. doi: 10.1002/bies.201300066.

34. Milo R, Jorgensen P, Moran U, Weber G, Springer M. 2010. BioNumbers--the database of key numbers in molecular and cell biology. Nucleic Acids Res. 38(Database issue):D750–3. doi: 10.1093/nar/gkp889.

35. Milo R, Phillips R. 2015. Cell Biology by the Numbers. 1st Edition. Garland Science. ISBN 9780815345374.

36. Moran MA et al., 2013. Sizing up metatranscriptomics. ISME J. 7(2):237–43. doi: 10.1038/ismej.2012.94.

37. Neurohr GE, Terry RL, Lengefeld J, Bonney M, Brittingham GP, Moretto F, Miettinen TP, Vaites LP, Soares LM, Paulo JA, Harper JW, Buratowski S, Manalis S, van Werven FJ, Holt LJ, Amon A. 2019. Excessive Cell Growth Causes Cytoplasm Dilution And Contributes to Senescence. Cell. 176(5):1083–1097.e18. doi: 10.1016/j.cell.2019.01.018.

38. Newman JR, Ghaemmaghami S, Ihmels J, Breslow DK, Noble M, DeRisi JL, Weissman JS. 2006. Single-cell proteomic analysis of S. cerevisiae reveals the architecture of biological noise. Nature. 441:840–6. doi: 10.1038/nature04785.

39. Neymotin B, Athanasiadou R, Gresham D. 2014. Determination of in vivo RNA kinetics using RATE-seq RNA. 20(10):1645–52. doi: 10.1261/rna.045104.114.

40. PaxDb: Protein Abundance Database. https://pax-db.org/

41. Pelechano V, Chávez S, Pérez-Ortín JE. 2010. A complete set of nascent transcription rates for yeast genes. PLoS One. 5(11):e15442. doi: 10.1371/journal.pone.0015442.

42. Pérez-Ortín JE, Alepuz P, Chávez S, Choder M. 2013. Eukaryotic mRNA decay: methodologies, pathways, and links to other stages of gene expression. J Mol Biol. 425(20):3750–3775. doi:10.1016/j.jmb.2013.02.029

43. Pérez-Ortín JE, Alepuz PM, Moreno J. 2007. Genomics and gene transcription kinetics in yeast. Trends Genet. 23(5):250–257.

44. Presnyak V, Alhusaini N, Chen YH, Martin S, Morris N, Kline N, Olson S, Weinberg D, Baker KE, Graveley BR, Coller J. 2015. Codon optimality is a major determinant of mRNA stability. Cell 160(6):1111–24. doi: 10.1016/j.cell.2015.02.029.

45. Radonjic M, Andrau JC, Lijnzaad P, Kemmeren P, Kockelkorn TT, van Leenen D, van Berkum NL, Holstege FC. 2005. Genome-wide analyses reveal RNA polymerase II located upstream of genes poised for rapid response upon S. cerevisiae stationary phase exit. Mol Cell. 18(2):171–183.

46. Raser JM, O’Shea EK. 2005. Noise in gene expression: origins, consequences, and control. Science. 309(5743):2010–2013.

47. Schönheit P, Buckel W, Martin WF. 2016. On the Origin of Heterotrophy. Trends Microbiol. 24(1):12–25. doi: 10.1016/j.tim.2015.10.003.

48. Schwanhäusser B, Busse D, Li N, Dittmar G, Schuchhardt J, Wolf J, Chen W, Selbach M. 2011. Global quantification of mammalian gene expression control. Nature. 473:337–342. doi: 10.1038/nature10098

49. Shah P, Ding Y, Niemczyk M, Kudla G, Plotkin JB. 2013. Rate-limiting steps in yeast protein translation. Cell. 153(7):1589–601. doi: 10.1016/j.cell.2013.05.049.

50. Siwiak M, Zielenkiewicz P. 2013. Transimulation - protein biosynthesis web service. PloS ONE. Sep 05 8(9): e73943 doi: 10.1371/journal.pone.0073943

51. Sun M, Schwalb B, Schulz D, Pirkl N, Etzold S, Larivière L, Maier KC, Seizl M, Tresch A, Cramer P. 2012. Comparative dynamic transcriptome analysis (cDTA) reveals mutual feedback between mRNA synthesis and degradation. Genome Res. J 22(7):1350–9. doi: 10.1101/gr.130161.111

52. Swanson M, Reguera G, Schaechter M, Neidhardt FC. 2016. Microbe, ASM Press. ISBN-10: 1555819125

53. Tani H, Akimitsu N. 2012. Genome-wide technology for determining RNA stability in mammalian cells: historical perspective and recent advantages based on modified nucleotide labeling. RNA Biol. 9(10):1233–8. doi: 10.4161/rna.22036.

54. Tarrant D, von der Haar T. 2014. Synonymous codons, ribosome speed, and eukaryotic gene expression regulation. Cell Mol Life Sci. 71(21):4195–206. doi: 10.1007/s00018-014-1684-2.

55. Velculescu VE, Madden SL, Zhang L, Lash AE, Yu J, Rago C, Lal A, Wang CJ, Beaudry GA, Ciriello KM, Cook BP, Dufault MR, Ferguson AT, Gao Y, He TC, Hermeking H, Hiraldo SK, Hwang PM, Lopez MA, Luderer HF, Mathews B, Petroziello JM, Polyak K, Zawel L, Kinzler KW, et al. 1999. Analysis of human transcriptomes. Nat Genet. 23(4):387–388.

56. von der Haar T. 2008. A quantitative estimation of the global translational activity in logarithmically growing yeast cells. BMC Syst Biol. 2:87. doi: 10.1186/1752-0509-2-87.

57. Wagner A. 2005. Energy constraints on the evolution of gene expression. Mol Biol Evol. 22(6):1365–1374. doi:10.1093/molbev/msi126.

58. Waldron C, Lacroute F. 1975. Effect of growth rate on the amounts of ribosomal and transfer ribonucleic acids in yeast. J Bacteriol. 122(3):855–865.

59. Warner JR. 1999. The economics of ribosome biosynthesis in yeast. Trends Biochem Sci. 24(11):437–440.

60. Wiechecki K, Manohar S, Silva G, Tchourine K, Jacob S, Valleriani A, Vogel C. 2017. Integrative metaanalysis reveals that most yeast proteins are very stable. bioRxiv 165290; doi: 10.1101/165290.

61. Yang E. et al., 2003. Decay rates of human mRNAs: correlation with functional characteristics and sequence attributes. Genome Res. 13(8):1863–72. doi:10.1101/gr.1272403

62. Zenklusen D.; D.R. Larson, R.H. Singer, 2008. Single-RNA counting reveals alternative modes of gene expression in yeast. Nat. Struct. Mol. Biol. 15: 1263–1271.

63. Zhang T, Wolfe C, Pierle A, Welle KA, Hryhorenko JR, Ghaemmaghami S. 2017. Proteome-wide modulation of degradation dynamics in response to growth arrest. Proc Natl Acad Sci U S A. 114(48):E10329–E10338. doi: 10.1073/pnas.1710238114.

